# OpenAntigens: a structure-aware database for antigen construct design across the human cell-surface and secreted proteome

**DOI:** 10.64898/2026.07.30.741735

**Authors:** Andre A. R. Teixeira, Haisun Zhu, Deepash Kothiwal, Ruili Cao, Allan Mills

## Abstract

Choosing which region of a protein to express remains poorly standardized in antibody discovery, recombinant reagent generation, structural biology and computational binder design. For human cell-surface and secreted proteins, this requires reconciling topology, processing, predicted and experimental structure, modifications, interaction partners, orthologs, paralogs and cross-reactivity risk before ordering DNA. OpenAntigens is a free, no-login database of construct-design reports for 5328 human secreted, GPI-anchored, single-pass and multipass proteins. It integrates UniProt topology, AlphaFold pLDDT and PAE, PDB precedent, InterPro and Pfam domains, mouse and cynomolgus orthologs, paralog and family context, Open Targets disease associations, partner and assembly context, and BLAST searches. It provides 55 305 construct suggestions spanning full design regions, PDB-backed boundaries, annotated domains, pLDDT/PAE-derived regions and membrane-expression options, plus 148 722 sequence-similarity hits to help choose constructs and assess cross-reactivity. For targets with compatible AlphaFold models, the interactive designer links sequence, structure, pLDDT and PAE, allowing users to revise boundaries and export species-equivalent sequences with real-time cysteine and modification warnings. OpenAntigens places reproducible construct suggestions, comparative context and browser editing in one workflow, reducing manual reconciliation across resources. OpenAntigens is available at openantigens.org.

**Graphical abstract:** 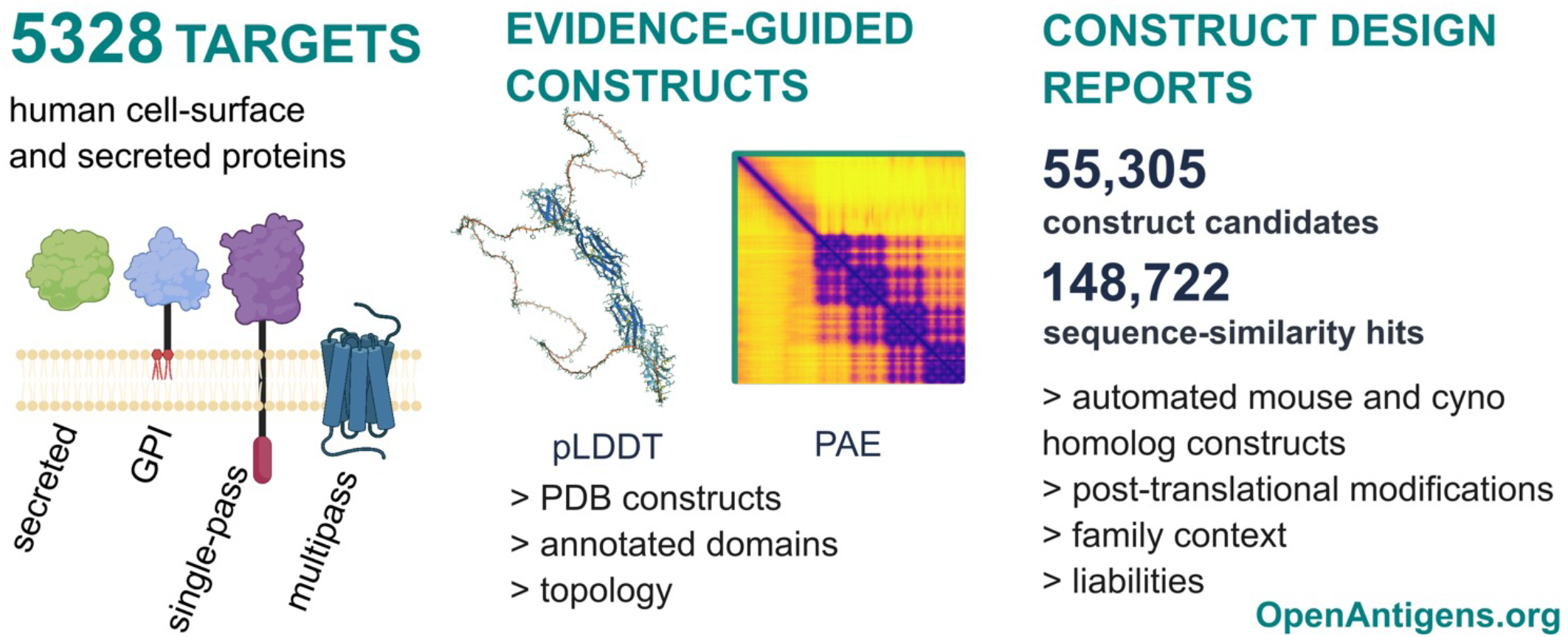

## Introduction

Recombinant target constructs are the physical entry point for many protein-science workflows: immunogens, panning antigens, biophysical reagents, structural-biology samples, assay standards and targets for computational binder design. The decision sounds simple: choose the region of a protein to express or display. In practice, it is a multi-source curation task. Users inspect UniProt for signal peptides, mature chains, topology, disulfides and modifications; AlphaFold DB for predicted structures, pLDDT and PAE; RCSB PDB for experimentally expressed boundaries; InterPro and Pfam for domains; ortholog resources for mouse or non-human-primate reagents; paralogs and BLAST for specificity risks; and disease or literature databases for prioritization [1-13]. These resources are powerful, but none is organized around the practical question of which construct to make.

The problem is particularly acute for human cell-surface and secreted proteins. Secreted proteins may need mature-chain boundaries and careful handling of propeptides, disulfides and cleavage sites. GPI-anchored and single-pass proteins usually need extracellular-domain constructs that keep conformational epitopes while excluding membrane and cytoplasmic regions. Multipass proteins may have short extracellular loops that are unsuitable as isolated soluble antigens, or large extracellular regions that need separate treatment from the membrane-spanning core. Some targets also require an interaction partner for efficient production or native presentation. Co-expression can rescue otherwise difficult proteins, and heteromeric systems need partner-chain assembly for correct folding. In each case, construct boundaries encode assumptions about folding, processing, assembly, cross-reactivity, surface exposure and specificity.

This target-side bottleneck matters more as AI-assisted binder design improves. Methods that design binders from target structures or surfaces have advanced rapidly [14-19], but they do not remove the need to decide which target, construct or epitope-containing region to use. A user still needs to know whether the chosen region is extracellular, structured, accessible, conserved in preclinical species when cross-reactivity is wanted, and divergent enough from paralogs when specificity is required. The same questions apply to in vitro display, immunization and hybrid workflows that combine computational design with experimental screening.

OpenAntigens addresses this gap by organizing heterogeneous public annotations and computed evidence into target-centric construct-design reports, focused on human proteins likely to be extracellularly accessible: secreted, GPI-anchored, single-pass and multipass proteins. For each target it resolves a canonical record, infers a design region, proposes construct candidates, maps ortholog-equivalent regions, summarizes paralog and family similarity, provides cross-reactivity context through local BLAST, reports sequence liabilities, flags assembly or partner-chain context, and presents disease and literature context. Each report is generated by the same pipeline, so users reviewing the same release start from the same evidence and algorithmically generated suggestions. OpenAntigens does not predict expression success or guarantee that a binder raised against a construct will be specific. Those outcomes depend on experimental factors the database does not model.

OpenAntigens draws on existing resources but addresses a construct-design use case that they do not cover in combination. Surfaceome and secretome resources establish which human proteins reach the cell surface or are secreted, including the SURFY in silico surfaceome [20], the mass spectrometry-based Cell Surface Protein Atlas [21], the Human Protein Atlas secretome [22] and MetazSecKB [23]. OpenAntigens uses this kind of evidence as one input to its target universe, but these resources stop at annotation and do not propose which region to express. Earlier tools support domain selection or domain-boundary prediction, including DomainView [24] and the high-throughput pipeline of Chen et al. [25]. They predate AlphaFold confidence metrics and do not combine surface or secreted scope with species-equivalent boundaries, paralog similarity and assembly context. OpenAntigens combines a defined surface and secreted target universe with AlphaFold pLDDT-and PAE-derived boundary logic, species-equivalent ortholog regions, paralog and BLAST context, partner and liability warnings, and in-browser construct export. By assembling these layers into one target-centric report per protein, OpenAntigens provides a single starting point for choosing surface and secreted antigen constructs, while the source databases remain authoritative for their respective annotations.

## Database content and construction

### Target universe and release snapshot

The OpenAntigens 2026-07-23 release covers 5328 human targets, selected from 6364 reviewed human Swiss-Prot proteins carrying a cell-membrane or secreted keyword after removing immunoglobulin and T-cell-receptor loci. Of these candidates, 5328 were retained because curated annotations supported cell-surface localization or secretion, and 1036 were excluded (Supplementary Methods). The release spans four topology buckets, secreted (1725), GPI-anchored (137), single-pass (1507) and multipass (1959) proteins. The release is designated 2026-07-23 based on its data-retrieval and processing date. The static portal, comprising target pages, JavaScript report data, flat-file downloads, license notes and release metadata, was generated on 2026-07-25 with software version 0.1.0. Figure 1 shows the construction workflow (a) and the release coverage across topology classes and evidence types (b-d).

**Figure 1.**
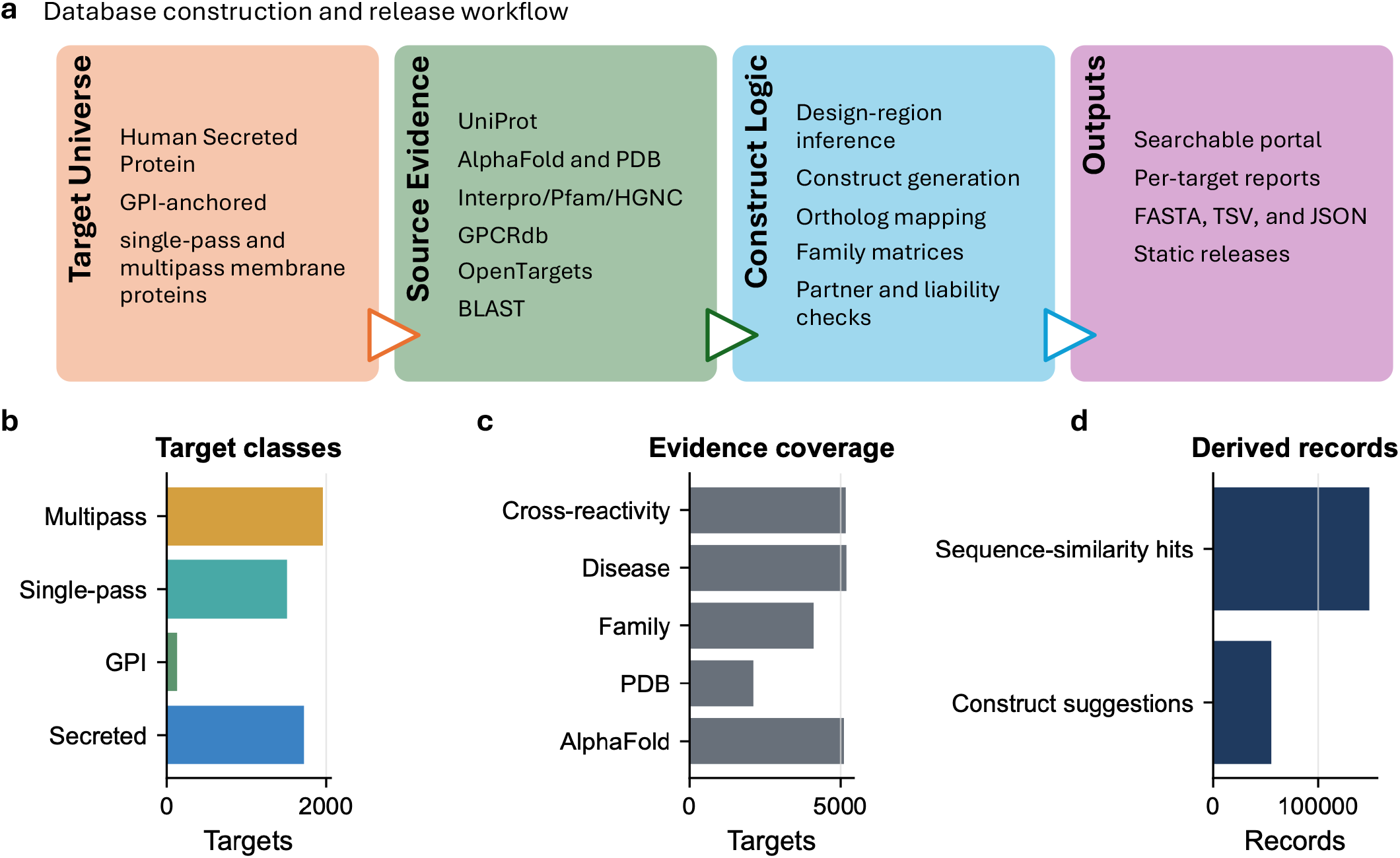
OpenAntigens organizes the human cell-surface and secreted proteome into structure-aware construct-design reports through a four-stage pipeline. (a) Build workflow: each target passes through a target universe (human secreted, GPI-anchored, single-pass and multipass membrane proteins), source evidence (UniProt, AlphaFold and PDB, InterPro/Pfam/HGNC, GPCRdb, Open Targets, BLAST), construct logic (design-region inference, construct generation, ortholog mapping, family matrices, partner and liability checks) and outputs (searchable portal, per-target reports, FASTA/TSV/JSON, static releases). (b) The 5328-target universe by membrane topology: 1959 multipass, 1725 secreted, 1507 single-pass and 137 GPI-anchored proteins. (c) Per-target evidence coverage, as targets out of 5328: AlphaFold models for 5107, a disease link for 5189, a cross-reactivity screen for 5167, family context for 4113 and an experimental PDB structure for 2114. (d) Records derived across the release: 55 305 construct suggestions and 148 722 sequence-similarity hits for cross-reactivity assessment. Almost every target carries a predicted structure and two in five carry an experimental one, so structure-aware design at this scale rests largely on AlphaFold. Candidate constructs and sequence-similarity hits are precomputed where the required sequence, annotation and structural evidence is available.

OpenAntigens is organized around targets rather than source databases. A user can search by UniProt entry name, accession, gene symbol, protein name, family label, topology class or disease context. Each target page then gathers the evidence needed for construct review: identity, aliases, topology, inferred design region, suggested constructs, AlphaFold and PDB evidence, domains, PTMs, cysteines, furin-like motifs, ligand annotations, assembly context, orthologs, family members, BLAST cross-reactivity, Open Targets associations and external links.

### Source data

OpenAntigens uses UniProt as the primary identity, sequence and feature source [1]. AlphaFold DB provides predicted structures, pLDDT and PAE when a compatible model exists [2,3]. RCSB PDB and wwPDB mappings provide experimental structure evidence and precedent boundaries [4,5]. InterPro and Pfam support domain and family annotation [6,7]; HGNC and Ensembl support gene nomenclature, family and paralog context [8,9]; NCBI RefSeq supports ortholog sequence context, particularly for cynomolgus monkey [10]; and GPCRdb provides GPCR segment, family and generic-numbering information [11]. UniProt interaction and subunit annotations and curated Complex Portal records support partner-chain and assembly warnings [1,26]. Open Targets provides disease-association context through direct and indirect association scores [12], and PubTator3 and PubMed links provide literature context [13]. Local BLAST searches against human, mouse and cynomolgus monkey protein databases provide cross-reactivity context [27]. Topology classification additionally draws on the SURFY in silico surfaceome [20] and the Human Protein Atlas secretome [22].

### Canonical mapping and design-region inference

OpenAntigens resolves each target to a canonical human protein record and records a warning where the mapping is ambiguous. The current release analyzes the canonical UniProt sequence for each target; alternative isoforms are not rendered as separate reports. Structure-dependent residue annotations are used only when the local structure sequence is compatible with the canonical sequence, because pLDDT, PAE, solvent accessibility, cysteine distances, topology labels and construct boundaries are all residue-indexed. Before generating candidates, OpenAntigens infers a design region: for secreted proteins, usually the mature chain after signal-peptide or propeptide removal; for GPI-anchored proteins, the extracellular region before the anchor signal; for single-pass proteins, the extracellular region defined by topology; and for multipass proteins, a distinction between large extracellular regions, which may receive soluble extracellular-region suggestions, and compact transmembrane proteins, treated primarily as membrane-expression targets.

### Construct recommendation workflow

OpenAntigens provides 55 305 construct suggestions across the release. These are reproducible hypotheses, not expression-success predictions, and the database deliberately shows five construct classes so users can choose between full-context, experimentally precedented, domain-focused, automatically calculated and membrane-expression strategies.

Full design-region constructs preserve the broadest inferred extracellular or mature-secreted context. They suit conformational epitopes that may span multiple domains, but may include flexible linkers, low-confidence segments or processing sites. PDB-backed constructs use experimental structure coverage mapped to UniProt coordinates and filtered to the design region. They provide precedent for a structurally characterized region, but do not establish ease of expression or purification, or reproduce all engineering used in the deposited construct. Domain-annotated constructs are derived from UniProt, InterPro and Pfam features within the design region, helping users select named modules such as immunoglobulin-like domains, receptor L-domains, cadherin repeats or growth-factor domains.

OpenAntigens also turns AlphaFold confidence into calculated candidates, running one procedure at two stringency settings. The lenient setting identifies regions with higher local confidence. It keeps residues above pLDDT 60, bridges short low-confidence gaps and leaves multi-domain units intact, returning the largest contiguous region supported by the model. The strict setting seeds from higher-confidence residues, above pLDDT 70, and then tests every internal boundary of a seed, asking whether each half is internally cohesive while their relative placement is uncertain. It cuts at the strongest boundary where the mean PAE between the two blocks is at least 12 and exceeds the PAE within them by at least 4, then repeats the same test within each resulting block, so a multi-domain seed is resolved recursively into its single domains rather than by one split (Supplementary Methods). Figure 2 shows both settings applied to one target. This uses pLDDT and PAE as construct-boundary evidence, beyond their usual role as visual diagnostics. A predicted structural unit is a candidate region for experimental testing; its boundaries do not establish independent folding after isolation from the surrounding sequence.

**Figure 2.**
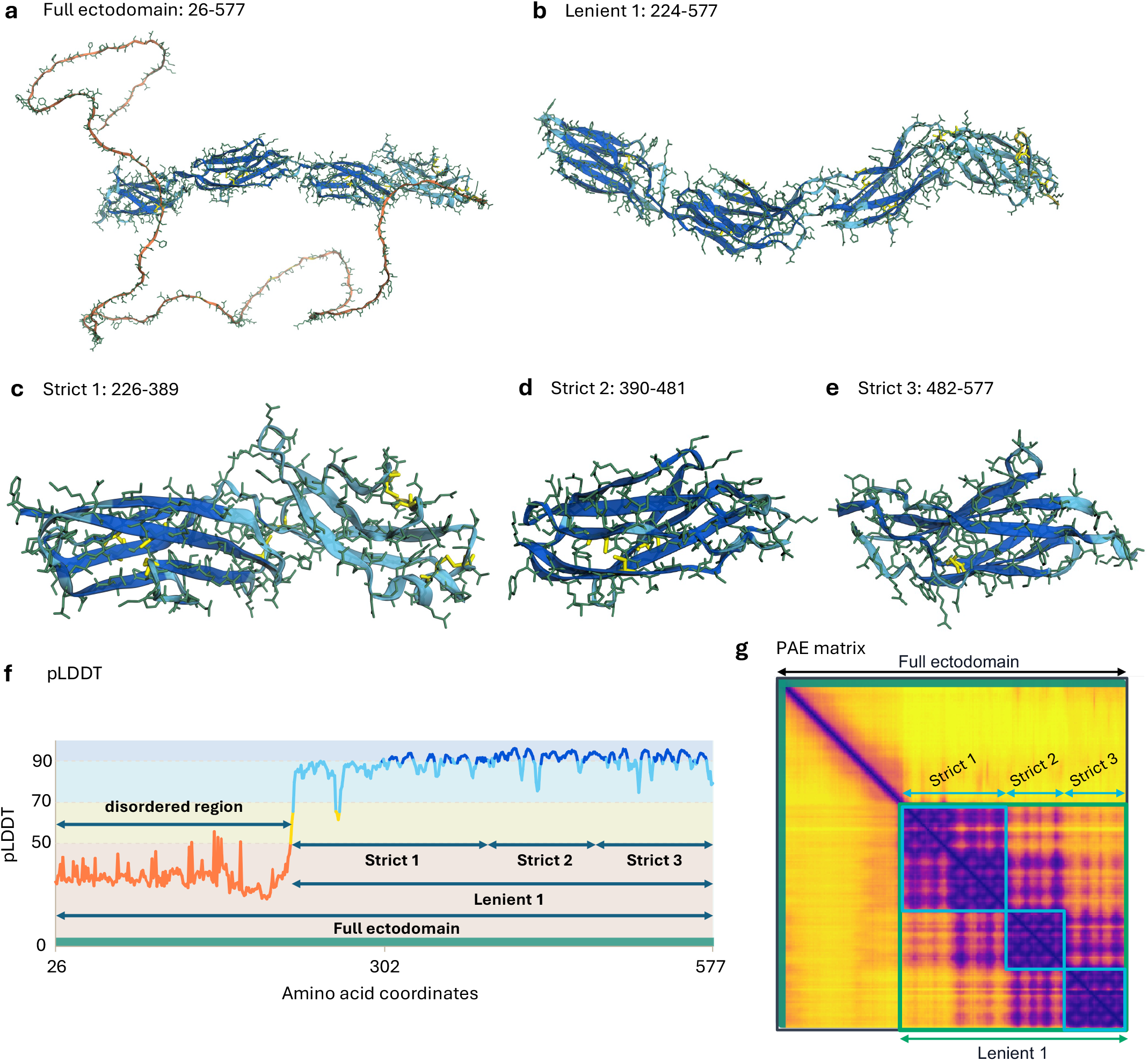
Per-residue confidence and predicted aligned error resolve the CD45 ectodomain into a nested set of designable constructs. (a-e) AlphaFold models of five suggested constructs, colored by per-residue pLDDT (blue high, orange low): (a) the full ectodomain (residues 26-577); (b) a lenient construct (224-577) that drops the low-confidence membrane-distal region; (c) strict construct 1 (226-389); (d) strict construct 2 (390-481); and (e) strict construct 3 (482-577). (f) Per-residue pLDDT along the ectodomain, with each construct drawn as a track beneath it. The membrane-distal region (about residues 26-220) falls below pLDDT 50 and overlaps a disordered region annotated in UniProt (28-163). (g) Predicted aligned error (PAE) matrix; dark on-diagonal blocks mark residue ranges whose relative geometry AlphaFold places with confidence, and cyan outlines mark strict constructs 1-3, the green outline marks the lenient construct (residues 224-577), and the black outline marks the full ectodomain (residues 26-577). The lenient construct removes the low-confidence N-terminal region, and the strict constructs divide the predicted structured remainder at PAE block boundaries into internally cohesive predicted units. Strict constructs 2 and 3 align with the two fibronectin type-III domains annotated in UniProt (391-483 and 484-576), recovering the domain architecture from the prediction alone. Captured from the live report at openantigens.org.

CD45 (PTPRC), a structured single-pass receptor-type tyrosine phosphatase, illustrates this logic (Figure 2). Its extracellular region (residues 26-577) contains a low-confidence membrane-distal segment followed by a series of predicted folded domains. The full design-region construct keeps the whole ectodomain; the lenient setting trims the low-confidence N-terminal segment; and the strict setting divides the predicted structured remainder at high-PAE junctions, two of which coincide with the protein’s fibronectin type-III modules. From one prediction a user can take a multi-domain construct for conformational epitopes that span domains, or an isolated domain for a focused reagent. CD45 carries 12 construct suggestions in all, spanning the full ectodomain, PDB-backed boundaries, annotated domains and both calculated settings.

Compact multipass proteins are handled separately. Their reports keep the target in its membrane context and emphasize topology, full-length cross-reactivity screens and optional membrane-engineering suggestions. These reports support selection of membrane-protein regions for expression, while researchers choose the expression format, detergents, lipids, stabilizing partners and assay conditions for their intended production system and assay.

For G-protein-coupled receptors, OpenAntigens also maps GPCRdb family, segment and generic-numbering annotations onto the sequence and structure, with residues labeled by extracellular, membrane or intracellular location. C-X-C chemokine receptor type 4 (CXCR4), a class A GPCR, illustrates what that adds (Figure 3). Its 352-residue chain forms a predicted seven-transmembrane bundle that the PAE matrix places within one internally cohesive predicted unit, so its construct suggestions span the full length. The labels then identify the N-terminus and three extracellular loops as segments that may be accessible to a cell-surface binder and provide the focus for cross-reactivity assessment.

**Figure 3.**
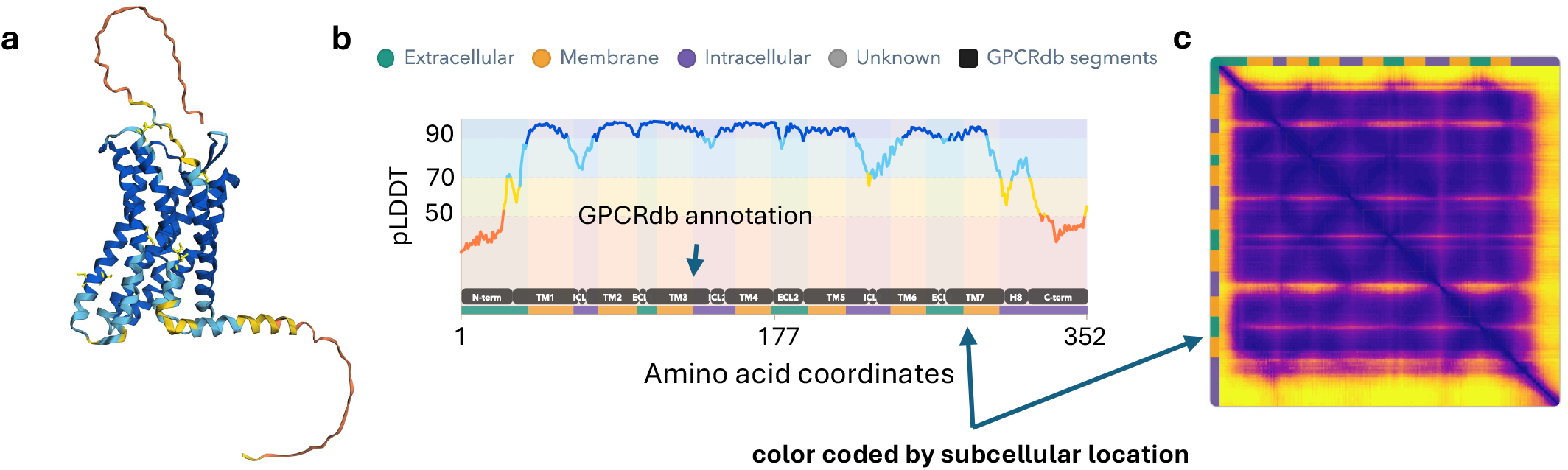
GPCRdb segment annotation maps every CXCR4 residue to a subcellular compartment and identifies extracellular segments that may be accessible to a cell-surface binder. (a) AlphaFold model colored by the predicted subcellular location of each segment. (b) Per-residue pLDDT across the 352-residue chain with the GPCRdb topology track beneath it: the N-terminus, seven transmembrane helices (TM1-TM7), the connecting intracellular and extracellular loops, helix 8 and the C-terminus, each colored by compartment (teal extracellular, orange membrane, purple intracellular, gray unknown). (c) Predicted aligned error (PAE) matrix. On a seven-transmembrane receptor, the potentially accessible target surface includes the N-terminus and three extracellular loops, which the GPCRdb track marks directly on the sequence. The PAE matrix places the helical bundle within one internally cohesive predicted unit, so construct suggestions span the full length while the extracellular segments provide the focus for accessibility and cross-reactivity assessment. Captured from the live report at openantigens.org.

Each construct card includes boundaries, sequence, score, classification, rationale, included domains, structural metrics, pLDDT and PAE diagnostics where available, PTMs, furin-like motifs, ligand annotations, cysteine warnings and ortholog-equivalent sequences. Cysteine warnings are construct-specific, flagging an unpaired cysteine or only one endpoint of a curated disulfide pair. PTM and processing annotations help users retain signal-processing context, keep disulfide-rich domains intact, and account for glycosylation and cleavage features relevant to expression or native presentation.

### Interaction partners and co-expression context

Many recombinant target failures are not solved by moving construct boundaries alone. Some proteins need stabilizing partners, obligate subunits or assembly context for efficient expression, secretion, trafficking or native antigen presentation. Co-expression can rescue production of a difficult protein, and at the extreme, heteromeric systems such as integrins require an alpha and beta chain for proper folding and presentation of the biologically relevant extracellular complex. OpenAntigens therefore includes assembly and interaction-partner context as a cautionary layer. When source annotations indicate a likely mandatory partner, obligatory complex or heterodimeric receptor, the report alerts the user that antigen design should proceed carefully and the primary literature should be reviewed before ordering constructs, preventing users from treating every target chain as an independent soluble antigen when native production may need a partner. The first release surfaces this known context so users can recognize when a target needs partner-chain design. Choosing partner boundaries, stoichiometry, vectors and linkers remains their decision. Explicit complex-aware antigen design, including partner-chain suggestions and co-expression planning, is planned for future releases.

### Homology, paralog and cross-reactivity context

Antigen design rarely stops at one human sequence. Users often want reagents that cross-react with mouse or cynomolgus monkey, or that avoid related human paralogs. OpenAntigens therefore treats orthology, paralogy and BLAST similarity as primary report content. For ortholog context, it maps the human design region to mouse and cynomolgus monkey sequences when suitable references are available, reporting construct-equivalent regions with sequence identity, coverage and extracted sequences, so a user can move from a human construct hypothesis to species-equivalent constructs without manually remapping boundaries. For paralog and family context, it combines InterPro family membership, Ensembl-derived paralog evidence and HGNC family information into family-member lists and directional identity and coverage matrices (Figure 4a). Directional metrics are retained because an extracellular region in one family member can be embedded in a larger region in another, making reciprocal comparisons non-identical. OpenAntigens also runs local BLAST screens against human, mouse and cynomolgus monkey protein sets, yielding 148 722 sequence-similarity hits across 5167 targets, with species, subject identifiers, E-value, bit score, identity, coverage and alignment coordinates. These are similarity signals, not pass/fail specificity predictions, that help users identify proteins to counter-screen, avoid, or consider as intended cross-reactive partners. Where compatible structures and annotations allow, reports emphasize residues that are both extracellular and surface accessible, computed from per-residue solvent accessibility [28,29] (Supplementary Methods), because antibodies and designed binders recognize accessible surfaces rather than all aligned residues equally. Figure 4 shows both layers for the UNC5 netrin-receptor family: the directional identity matrix across all five human members (Figure 4a), and, for UNC5B, an alignment against its closest human paralog UNC5C, with solvent-accessible UNC5B residues highlighted and identical positions marked between the rows (Figure 4b). The overlap between sequence identity and predicted surface accessibility identifies conserved surface positions that may warrant counter-screening. UNC5B is a typical case, with 15 construct suggestions and 49 similarity hits to weigh when deciding which relatives are acceptable cross-reactants and which to counter-screen before reagent generation.

**Figure 4.**
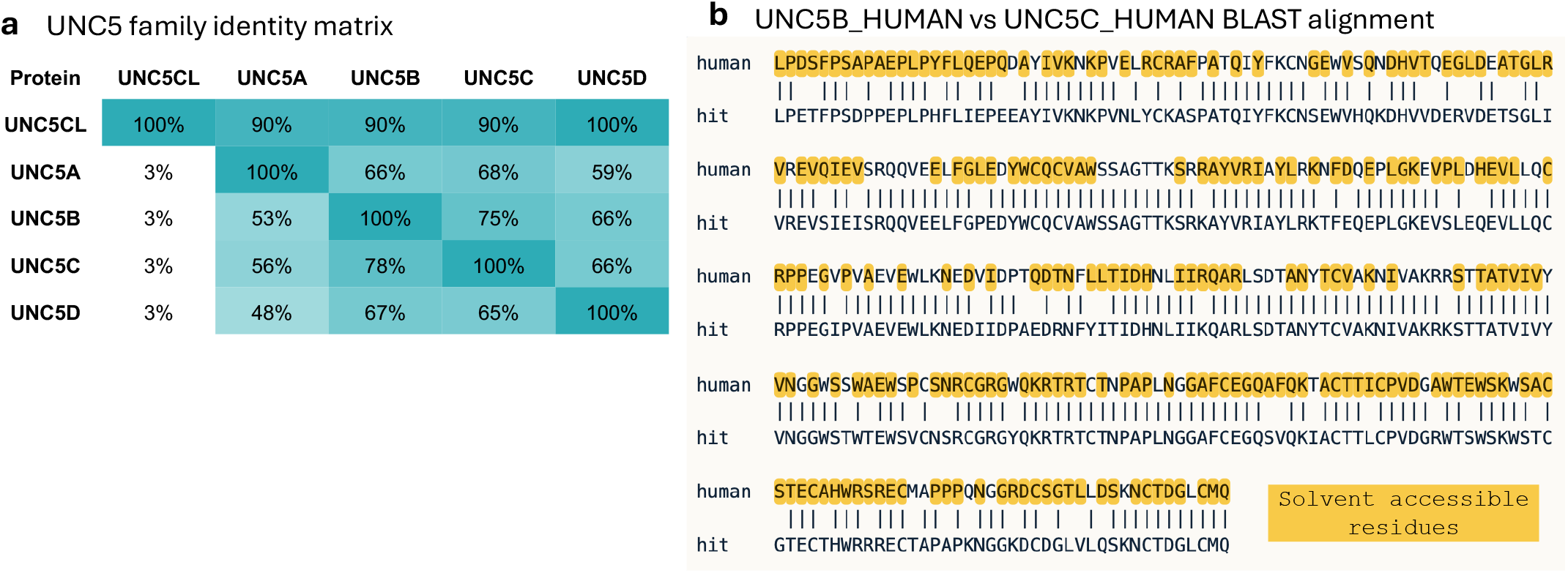
Directional family identity and a solvent-accessible paralog alignment show which conserved residues lie on the accessible surface. (a) Directional identity matrix across the five human UNC5 family members (UNC5CL, UNC5A, UNC5B, UNC5C, UNC5D). Each cell reports identical aligned residues as a percentage of the row protein’s ectodomain length, so the matrix is asymmetric. UNC5CL has a 10-residue ectodomain, so its row reads 90-100% against every paralog (top row) while those longer paralogs share about 3% of their own ectodomain with it (left column), which is why a percent identity is meaningful only once you say which protein it is a fraction of. (b) BLAST alignment of human UNC5B against its closest human paralog UNC5C, the highest-ranked human hit in its similarity screen. Identical positions are marked between the rows, and solvent-accessible residues in the UNC5B predicted structure are highlighted. Positions that are both identical and predicted to be accessible identify conserved surface regions that may warrant counter-screening. The matrix in (a) is redrawn from the live report data and reproduces its values. Panel (b) is a direct screen capture from the live report at openantigens.org.

### Web interface and downloads

The public portal at openantigens.org is static, free to use and requires no login. The index page provides global search, a disease-focused filter, sortable topology and structure-evidence columns, and links to target reports. Each report organizes target identity, design region, construct suggestions, structural support, domain architecture, sequence liabilities, homology, family context, cross-reactivity, disease associations and external resources. Construct Details are grouped into tabs by candidate type, and a Sections menu provides direct navigation through each report. The construct-selection guide describes how to compare candidates and inspect termini, cysteines and processing features. Guidance linked beside sequence export identifies the subsequent choices of expression host, vector, secretion leader and tags.

### Interactive construct designer

Reports with a compatible local AlphaFold model embed an interactive construct designer so precomputed suggestions, such as the CD45 constructs in Figure 2, can be inspected and adjusted in the browser without writing code. The designer assists with selecting and refining target protein regions for expression; selection of expression hosts, signal peptides, affinity tags, vectors and regulatory elements remains part of the user’s experimental design. The designer links a residue-level sequence view, a three-dimensional AlphaFold model colored by pLDDT, an interactive pLDDT trace and an interactive PAE matrix. Selecting a residue or contiguous range in any panel highlights the same window in the others, so a candidate boundary can be reviewed against folded domains, linkers and disulfides without reconciling coordinates across separate tools. Clicking a precomputed construct loads its boundaries into the same workspace. Reports without a compatible local AlphaFold model omit the interactive construct designer but retain the remaining target annotations and any precomputed construct information available for that target.

For the active selection, the designer produces export-ready records: the human construct name in GENE_SPECIES_start-end format, its boundaries and sequence, and equivalent mouse and cynomolgus monkey regions with external links where ortholog mappings exist. Identity to the human sequence is reported across the full selection and, where compatible structures and annotations permit, across residues classified as both extracellular and surface accessible. Records can be copied in TSV or FASTA format.

The designer also applies construct-level liability checks in real time. It flags unpaired cysteines, including cases where only one partner of a curated disulfide falls inside the selection; lists post-translational modifications, processing features and furin-like motifs contained in the selection; and offers optional cysteine-to-serine edits that update both the exported sequence and construct name. A selected region can be isolated in the structure view, and users can download either the full model or the selected region as a PDB file.

### Downloads and release snapshots

OpenAntigens provides flat-file downloads comprising a compact portal index in TSV and JSON and Open Targets disease-association data in TSV and JSON, with a manifest recording file descriptions, columns and source notes. The index includes entry name, gene symbol, protein name, topology bucket, family name, top disease, disease count, status, construct count, cross-reactivity count, structure-evidence flags, family-context flags and report links. Target reports are rendered as static HTML with per-target JavaScript payloads for interactive features, and the archived release includes a corresponding JSON record for each target. Each target has a stable report URL. The 2026-07-23 release is archived on Zenodo, and future dated releases will be archived separately (Supplementary Methods). Release metadata record the database name, software version, build and generation dates, target, status and topology counts, and licensing. Dated archives preserve the exact released content while the live site is refreshed as upstream resources change. The project will be maintained for at least five years after publication, with approximately six-month public data refreshes and dated releases planned.

## Discussion

OpenAntigens addresses a practical bottleneck in protein reagent generation: collecting and reconciling scattered target evidence before choosing constructs. By organizing topology, structure confidence, domain annotations, PDB precedent, orthologs, paralogs, BLAST similarity, PTMs, cysteines, disease context and literature links into target-level reports, the database makes construct triage more systematic and auditable. The same content supports two uses: a downloadable catalog of per-target construct evidence for programmatic pipelines, and an interactive per-target workspace for choosing and exporting constructs by hand.

The resource suits both classical and emerging workflows. Antibody-discovery teams can select immunogens, panning targets, recombinant screening antigens and counter-screening proteins. Protein biochemists can compare expression constructs, identify sequence liabilities and recognize targets that may need co-expression with a partner. Structural biologists can use PDB precedent and PAE-guided regions as construct starting points. Computational binder-design users can define extracellular target constructs, identify conserved species-reactive surfaces and avoid paralog-conserved regions before launching design calculations.

Practical resources can help researchers plan recombinant protein expression after selecting a target region. The protein production and purification guide from the Structural Genomics Consortium and collaborating centers provides a general starting strategy [30]. Detailed protocols describe expression in *Escherichia coli* [31], Expi293F cells [32], ExpiCHO and HEK293E cells [33], and baculovirus-infected insect cells [34]. Guides to affinity tags [35] and recombinant protein purification [36] can help researchers choose tags and plan purification workflows. These resources are linked in the website’s After export guidance.

Several limitations should guide interpretation. Predicted structures and confidence metrics are not experimental validation. AlphaFold pLDDT and PAE can help identify plausible structured regions and domain boundaries, but they do not guarantee expression, folding, stability, secretion or binder compatibility. The boundary-selection procedure does not use secondary-structure assignments to constrain cuts to loops or exclude helices and strands. Users should inspect both termini in the structure view and refine them where needed before experimental testing. Of 5328 reports, 221 lack a compatible AlphaFold DB structure incorporated into the release and therefore omit AlphaFold-dependent residue calculations and interactive structure views. The current release analyzes only the canonical UniProt sequence, so alternative isoforms and isoform-specific topology may not be represented. OpenAntigens also inherits limitations from upstream annotations: missing topology, incomplete PTM records, missing partner annotations and incomplete ortholog references can affect report completeness. BLAST similarity provides cross-reactivity context but is not an antibody-binding assay. Multipass membrane proteins and protein complexes remain difficult, and automated suggestions should be reviewed in the context of the expression system, membrane environment, co-expression strategy, assay format and biological state.

We plan to incorporate additional structure prediction models into OpenAntigens, prioritizing proteins without compatible AlphaFold DB structures. The CD45 comparison illustrates why the pLDDT and PAE thresholds used for construct design should be evaluated before adding another predictor (Supplementary Figure 1). Although the models show similar organization of the folded domains, ESMFold2 assigns higher pLDDT to the N-terminal region annotated as disordered in UniProt (mean 66.7 versus 34.4 for AlphaFold), so the default lenient setting retains this region. Lower PAE between the two C-terminal regions also prevents the default strict setting from separating them. Calibration should therefore assess each predictor across multiple proteins, using curated domain boundaries and experimental construct performance to evaluate the resulting designs.

Future releases will improve design support for obligate protein complexes, particularly systems such as integrins that require partner-chain context. Where feasible, they will also add predicted structures for targets that lack a compatible AlphaFold DB model, subject to practical limits for very long proteins. An API and Model Context Protocol (MCP) interface are planned to support automated queries and interactions with OpenAntigens records. A longer-term goal is to connect these annotations to binder-design engines so users can select conserved or specificity-favoring target regions and launch design calculations in the same biological context. We also plan to extend coverage to intracellular proteins and evaluate support for user-supplied sequences, with appropriate handling of topology and processing and the computation required for new analyses.

## Acknowledgements

We thank Dr. Kenneth Fasman for substantive review and suggestions on the manuscript and for his ongoing advice and support throughout the project. We thank Hoan Nguyen, Yang Su and Kunal Shah for testing the OpenAntigens website and proposing improvements, changes and fixes. We also thank the teams that create and maintain the public databases, datasets and open-source software used by OpenAntigens, including UniProt, AlphaFold DB, RCSB PDB, InterPro, Pfam, HGNC, Ensembl, NCBI, GPCRdb, Complex Portal, Open Targets, PubTator3, SURFY, the Cell Surface Protein Atlas, the Human Protein Atlas, MetazSecKB, BLAST, FreeSASA and 3Dmol.js.

## Author contributions

Andre A. R. Teixeira: Conceptualization, Methodology, Software, Data curation, Formal analysis, Validation, Visualization, Project administration, Writing - original draft, Writing - review and editing. Haisun Zhu, Deepash Kothiwal, Allan Mills and Ruili Cao: Conceptualization, Methodology, Software, Validation, Writing - review and editing.

## Supplementary data

Supplementary Data are available at NAR Online. The supplementary file contains Supplementary Methods and Supplementary Figure 1, which compares the CD45 construct boundaries obtained from AlphaFold and ESMFold2 predictions. Detailed methods for the release build, topology and structure processing, construct generation, partner warnings, cross-reactivity analysis and download generation are provided in the Supplementary Methods.

## Conflict of interest

The authors declare no competing interests.

## Funding

This work was made possible by the generosity of Tim Springer, Chafen Lu and their family, whose gift established the endowment that supports research at the Institute for Protein Innovation. Funding for open access charge: Institute for Protein Innovation.

## Data availability

OpenAntigens is publicly available at openantigens.org. The public site provides static target pages and downloadable TSV/JSON files. The current download manifest includes a compact portal index in TSV and JSON format and Open Targets disease-association files in TSV and JSON format. OpenAntigens-generated annotations are distributed under the Creative Commons Attribution 4.0 license (CC BY 4.0) unless otherwise noted. Third-party source data retain their original licenses, terms of use and citation requirements. The live portal is periodically refreshed. The exact 2026-07-23 release described here is archived as a frozen snapshot at Zenodo (https://doi.org/10.5281/zenodo.21628751), which is the version of record for the numbers reported in this paper.

## Code availability

The OpenAntigens software is released under the Apache License 2.0 at https://github.com/proteininnovation/OpenAntigens. The repository includes the analysis pipeline, static portal renderer, release scripts, documentation and tests. The data and rendered portal for the 2026-07-23 release are archived in the Zenodo deposit cited above.

## Supplementary Information for OpenAntigens

Supplementary Methods and Supplementary Figure 1 for *OpenAntigens: a structure-aware database for antigen construct design across the human cell-surface and secreted proteome*. Reference numbers in square brackets refer to the reference list of the main article.

## Supplementary Methods

### Release build and target processing

The target universe was defined from reviewed human UniProtKB/Swiss-Prot entries carrying a Cell membrane or Secreted keyword, after removing immunoglobulin and T-cell-receptor loci, giving 6364 candidate proteins. Each candidate was assigned a primary topology class by parsing UniProt subcellular-location and topology annotations (secreted soluble, single-pass surface, GPI-anchored surface, multipass surface, mixed secreted and surface, surface-topology-unspecified, or isoform-specific surface or secreted), with the SURFY in silico surfaceome [20] and the Human Protein Atlas and Uhlen secretome tables [22] used as orthogonal support and as a rescue layer for ambiguous cases. Proteins with qualifying annotations supporting secretion or cell-surface localization were retained, including secreted, single-pass, GPI-anchored, multipass, mixed and isoform-specific cases. Peripheral-membrane-associated proteins and topology-unspecified membrane proteins without surfaceome support were excluded. This produced 5328 retained targets and 1036 exclusions. Retained targets are reported in four topology buckets (secreted, GPI-anchored, single-pass and multipass) and tagged with an accessibility-confidence tier (high, medium, isoform-specific or therapeutic-rescue); a small number enter a bucket through isoform-specific annotation rather than canonical topology, so the bucket counts are curated retained-target counts, not pure topology classes. Each retained target was resolved to a human protein record, analyzed through the OpenAntigens pipeline and rendered into a static portal snapshot. The 2026-07-23 release snapshot contains 5328 target rows, 5328 successful reports and no error records. Batch-level metadata, topology counts and status counts were read from the release portal_metadata.json and batch_summary.json files.

### Topology, structure and domain processing

Topology and design regions were inferred from UniProt signal peptide, chain, topological domain, transmembrane, propeptide and anchor-related features. All sequence-level analyses use the canonical UniProt sequence; alternative isoforms are not analyzed as separate targets. AlphaFold DB structures and PAE files were retrieved or reused from local cache when available and were accepted for residue-level calculations only when sequence-compatible with the canonical target. Compatible AlphaFold structure and PAE files were incorporated for 5107 of 5328 targets. The remaining 221 reports omit AlphaFold-dependent residue calculations and interactive structure views, although 95 retain experimental PDB evidence and 85 retain one or more precomputed construct suggestions. pLDDT values were used for local-confidence tracks and PAE matrices were used to evaluate relative confidence between candidate regions. PDB-backed construct candidates were derived from experimental structure coverage mapped to UniProt coordinates and filtered against the target design region. Domain candidates were derived from UniProt, InterPro and Pfam annotations after filtering for design-region relevance.

### Construct generation and annotation

OpenAntigens generated construct candidates from full design regions, PDB-backed boundaries, domain annotations, lenient pLDDT-supported regions, strict pLDDT and PAE-supported regions, and membrane-expression tracks where appropriate. The lenient setting retained higher-confidence segments at a per-residue pLDDT threshold of 60, bridged low-confidence gaps of up to 12 residues, and kept larger multi-domain units without further splitting. The strict setting seeded segments at pLDDT 70, bridging low-confidence gaps of up to 8 residues, then split each seed into single domains using PAE. At each recursion level, every cut that left at least 40 residues on each side was tested. A cut was eligible when mean inter-block PAE was at least 12 and exceeded mean intra-block PAE by at least 4. Eligible cuts were ranked by separation + 0.35 x max(0, local boundary PAE - intra-block PAE) + linker bonus; local boundary PAE and linker pLDDT were calculated over windows extending up to five residues on each side of the cut, and the linker bonus increased as mean linker pLDDT fell below 70. The top-ranked cut was applied recursively, to a maximum depth of four. Both settings required segments of at least 25 residues; all constructs were then filtered to a minimum length of 50 residues and to the inferred design-region boundaries. Exact duplicate boundaries among calculated and annotated candidates were collapsed, retaining the higher-scoring candidate. PDB-backed candidates were exempt and retained individually even when their coordinates matched another construct, because each deposited structure provides separate experimental evidence. The full-length membrane-expression construct was also retained as a separate baseline. Construct cards were annotated with sequence, boundaries, classification, rationale, included domain annotations, structural metrics, PTMs, cysteine context, furin-like motifs, ligand annotations, homolog-equivalent sequences and warnings. Exact termini follow the retained pLDDT intervals and any accepted PAE cuts, subject to the design-region and length filters above. The procedure does not use secondary-structure assignments to constrain cuts to loops or exclude helices and strands. Users should inspect both termini in the structure view and refine them where needed before experimental testing.

### CD45 comparison with ESMFold2

For Supplementary Figure 1, the CD45 ectodomain (PTPRC, UniProt P08575, residues 26-577) was analyzed using the full-length AlphaFold DB structure and PAE matrix from the 2026-07-23 release snapshot and a supplied ESMFold2 prediction of the ectodomain. The ESMFold2 sequence matched canonical residues 26-577 exactly; model positions 1-552 were mapped to those coordinates. One prediction per method was analyzed using the construct-generation rules above without changing any parameters. Structure views were aligned with PyMOL using C-alpha atoms in canonical residues 233-576.

**Supplementary Figure 1.**
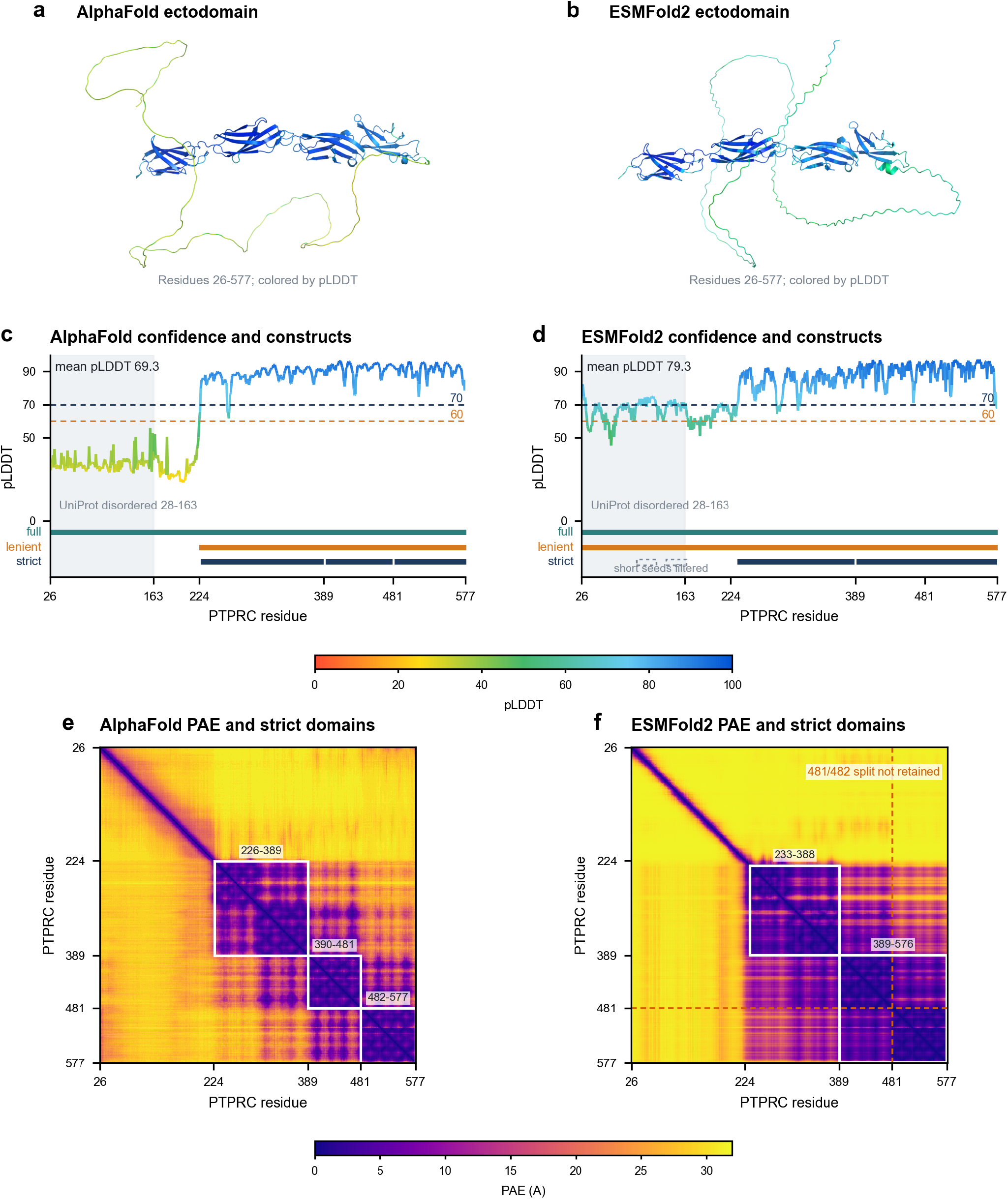
AlphaFold and ESMFold2 confidence estimates yield different CD45 construct boundaries under the same design parameters. (a, b) PTPRC/CD45 ectodomain structures (UniProt P08575, residues 26-577), from the full-length AlphaFold DB model and an ESMFold2 prediction of the ectodomain sequence, respectively. Structures are shown in a common orientation after superposition of the folded region and colored by predicted local distance difference test (pLDDT) score. (c, d) Per-residue pLDDT and construct intervals calculated with the OpenAntigens website defaults. Gray shading denotes the UniProt disordered region (28-163); horizontal dashed lines mark the lenient (60) and strict (70) pLDDT thresholds. Teal, orange and navy tracks indicate the full ectodomain, lenient intervals and retained strict constructs, respectively. The ESMFold2 lenient interval (26-577) duplicates the full ectodomain and is removed during deduplication. Gray dashed boxes mark strict seeds excluded by the 50-residue minimum construct length. (e, f) Predicted aligned error (PAE, angstroms) matrices, with white boxes marking retained strict constructs. Orange dashed lines in (f) mark the AlphaFold boundary at 481/482, which does not meet the default PAE splitting criteria for the ESMFold2 prediction. Both predictions were analyzed with identical thresholds and filtering rules (Supplementary Methods); all coordinates refer to canonical PTPRC residues.

### Interaction-partner and assembly warnings

OpenAntigens extracted assembly and partner context from source annotations including UniProt subunit and interaction records, Complex Portal records where available, and curated target-class logic for known heteromeric systems. UniProt SUBUNIT annotations and curated Complex Portal records provide interaction and assembly context. Human Complex Portal records are present for 630 reports in this release. Automated obligatory-partner warnings are limited to 26 reports for canonical integrin alpha or beta chains and use curated family logic together with UniProt SUBUNIT text and Complex Portal records where available. Other interaction or complex-membership records are presented as context and do not by themselves establish a requirement for co-expression. The current release does not design multi-chain constructs. The pipeline does not reconstruct biological homo-oligomeric assemblies or incorporate interfaces between their subunits into the selection of construct boundaries.

### Ortholog, paralog and cross-reactivity analysis

Ortholog reference sequences were precomputed from the HGNC Comparison of Orthology Predictions (HCOP) [8]. For each canonical reviewed human UniProt target, HCOP was queried for mouse (taxon 10090) and cynomolgus monkey (Macaca fascicularis; taxon 9541) orthologs. When HCOP returned multiple predictions, mouse calls were ranked by gene-symbol source (MGI, then VGNC, then NCBI) and cynomolgus calls by VGNC then NCBI; the number of supporting HCOP sources was the next ranking criterion. Mouse proteins were resolved by the HCOP gene symbol, with the human gene symbol as a fallback lookup key, against a local reviewed UniProtKB/Swiss-Prot proteome. An unreviewed sequence from mouse reference proteome UP000000589 was used only when no reviewed protein was available for that gene. Cynomolgus proteins were resolved by NCBI Gene ID, HCOP gene symbol or, as a final lookup key, the human gene symbol against the annotated protein set of RefSeq assembly GCF_037993035.2 [10]. NP_ products were preferred to XP_ products, and the longer product was selected among candidates of the same class. The release resolved mouse sequences for 5004 of 5328 targets and cynomolgus sequences for 4593 targets; unresolved species mappings are reported as unavailable.

The complete human and ortholog sequences were globally aligned to project the human design-region coordinates onto each ortholog. Global alignments used a match score of 1, a mismatch score of 0 and a unit gap penalty. The projected ortholog region was then aligned to the human design region to calculate identity and coverage. Each human construct was mapped through this design-region alignment. Its ortholog coordinates span the first to last mapped ortholog residue, and any human residues aligned to gaps are recorded in the mapping notes. The report and interactive designer provide the ortholog accession, projected coordinates, sequence, full-selection identity and, when structural annotations permit, extracellular accessible identity. These records are also included in construct-level TSV and FASTA exports.

Canonical family assignment used InterPro records explicitly annotated as families [6]. When several InterPro family records were available, candidates were ranked using sequence coverage, the number of annotated fragments and similarity between the family and target names. Precomputed family sets grouped proteins in the OpenAntigens target universe that shared the selected InterPro accession and compared their inferred extracellular design-region sequences. These InterPro-derived sets were the primary family and paralog source. Ensembl human paralogs [9] and HGNC family members [8] were queried only when no precomputed InterPro family members were available. Candidate proteins were resolved to human UniProt sequences and retained only when a design region could be derived. At report generation, OpenAntigens first loaded the precomputed paralog set, followed by the precomputed InterPro family alignment. Live HGNC family context was used only for targets without a usable precomputed InterPro family assignment. Detailed contexts containing more than 20 members were not loaded. Directional identity from row protein i to column protein j was calculated as 100 times the number of identical aligned residues divided by the length of the row protein’s design region. Directional coverage was 100 times the number of aligned, nongap row residues divided by the same length. The row-specific denominators allow reciprocal values to differ. For multipass mixed cases and multipass targets without an extracellular design region, an additional full-length family context was generated when the required family-member sequences were available.

Sequence similarity was calculated with local blastp searches [27]. Reports with an inferred extracellular design region searched that region. Multipass targets and targets without such a region were also searched using the full canonical sequence. The primary databases contained reviewed UniProt human, mouse and cynomolgus monkey proteins. If a query returned no hit from the reviewed cynomolgus set, it was searched against a broader local cynomolgus UniProt protein set. Searches used an E-value threshold of 1 × 10^-5 and requested at most 25 target sequences per species. Hits containing the query accession or UniProt entry name in the subject identifier were removed. Duplicate subject identifiers were reduced to the hit with the highest bit score, followed by coverage and identity, and the retained hits were ordered by decreasing bit score and increasing E-value.

Cynomolgus BLAST hits underwent an additional reporting step. Hits were grouped by gene symbol, and the hit with the highest bit score, coverage and identity was selected for each gene. That gene was resolved to a RefSeq protein, and identity, coverage, coordinates and aligned sequences were recalculated by global alignment to the RefSeq sequence. Only successfully remapped cynomolgus hits were retained. The bit score and E-value from the initial BLAST hit were retained for ranking.

For reports with a compatible AlphaFold structure and PAE matrix, per-residue solvent-accessible surface area was calculated with FreeSASA [28] using the Lee-Richards algorithm [29], a 1.4-angstrom probe radius and 20 slices. Relative SASA was calculated from residue-specific maximum accessible areas. A residue was classified as exposed when its full-model relative SASA was at least 0.20. Residues below that threshold were recalculated within a selected structural block, using strict pLDDT/PAE-derived regions when available and topology-derived regions otherwise; residues meeting the 0.20 threshold in this context were classified as conditional. SASA was not evaluated for residues with pLDDT below 50. When represented in the structure, these residues were classified as conditional rather than buried. Both exposed and conditional classes were included in the extracellular accessible residue set. Extracellular accessible identity was the fraction of aligned, nongap primary-target positions that were identical to the comparison sequence among positions classified as both extracellular and accessible. This target-directed measure was added to ortholog, family and BLAST comparisons and to construct exports when the required structure, topology and alignment data were available. These calculations use the target-chain model and selected structural blocks without partner chains. Residues classified as exposed may be buried against another subunit, including an identical copy, in the biological assembly.

### Disease, literature and download generation

Open Targets disease associations were built from bulk Open Targets release files by joining target, disease and association tables locally. The portal index displays top disease context and target pages include direct and indirect association scores when available. PubTator3 links and counts provide literature context. The static portal builder generated report pages, JavaScript report payloads, a compact TSV/JSON portal index, Open Targets TSV/JSON files, a download manifest and release metadata. Each of the 5107 reports with a compatible AlphaFold model embeds a client-side interactive construct designer that renders the model with 3Dmol.js and links the sequence, structure, pLDDT and PAE views through one residue selection. The remaining 221 reports omit the interactive designer but retain the other target annotations and any precomputed construct or PDB evidence available for that target.

Target records are addressed by UniProt entry name. In the live portal, PTPRC_HUMAN resolves to reports/ptprc_human.html and loads its report data from a JavaScript payload. In the Zenodo machine-readable archive, the corresponding JSON record is reports/human/ptprc_human_report.json. Each dated release is archived with its own DOI.

### Use of AI tools

The authors conceived, designed and implemented OpenAntigens, including its pipeline, analyses and scientific conclusions. Large language models (OpenAI GPT-5.4, GPT-5.5 and GPT-5.6; Anthropic Claude Opus 4.7, 4.8 and 5) were used to assist software development and code review and to review and edit the manuscript. AI-assisted code was validated through the project’s automated test suite, including golden-output regression tests, and by checking pipeline outputs against the underlying source records. The authors take responsibility for the content of this work.

## Notes

https://openantigens.org/

https://github.com/proteininnovation/OpenAntigens

https://doi.org/10.5281/zenodo.21628751

### Competing Interest Statement

The authors have declared no competing interest.

### Summary of Updates

Revised the manuscript to clarify construct selection, ortholog mapping, and limitations of structure-based recommendations. Expanded the Supplementary Methods and added a comparison of AlphaFold and ESMFold2 predictions for CD45 in Supplementary Figure 1. Updated Figure 2 and its legend, clarified interaction-partner and assembly context, and expanded the Discussion with practical guidance and references for recombinant protein expression and purification. Reformatted the manuscript and incorporated the supplementary information at the end of the PDF.

